# Image-Based Recognition of Parasitoid Wasps Using Advanced Neural Networks

**DOI:** 10.1101/2024.01.01.573817

**Authors:** Hossein Shirali, Jeremy Hübner, Robin Both, Michael Raupach, Stefan Schmidt, Christian Pylatiuk

## Abstract

Hymenoptera have some of the highest diversity and number of individuals among insects. Many of these species potentially play key roles as food sources, pest controllers, and pollinators. However, little is known about their diversity and biology, and about 80% of the species have not been described yet. Classical taxonomy based on morphology is a rather slow process, but DNA barcoding has already brought considerable progress in identification. Innovative methods such as image-based identification and automation can even further speed up the process. We present a proof of concept for image data recognition of a parasitic wasp family, the Diapriidae (Hymenoptera), obtained as part of the GBOL III project. These tiny (1.2 - 4.5 mm) wasps were photographed and identified using DNA barcoding to provide a solid ground truth for training a neural network. Subsequently, three different neural network architectures were trained, evaluated, and optimized. As a result, 11 different classes of diaprids and one class of “other Hymenoptera ’’ can be classified with an average accuracy of 96%. Additionally, the sex of the specimen can be classified automatically with an accuracy of > 96%.

## 1. Introduction

While it is common knowledge that the highest (insect) diversity is found in the tropics (Godfray et al. 1999, Dunn and Fitzpatrick 2012), several recent studies (e.g., Chimeno et al. 2022, 2023) suggest there is even in Germany a huge number of unknown arthropod species. The majority of those taxa are among the insect domains Diptera and Hymenoptera, and they are referred to as “dark taxa” (Hartop 2021). Another common denominator is their small body size: the highest diversity and individual numbers among insects can be found in the small-bodied groups (but not because of their size (Rainford et al. 2016)), making even basic tasks such as specimen handling and mounting a challenge (Morinière et al. 2019). Although many of these species play potentially key roles in all kinds of habitats as food sources, pest controllers, pollinators, etc., little is known about their diversity and biology (Dunn and Fitzpatrick 2012).

Hallmann et al. (2017) recorded a devastating 75% decline in insect biomass within 27 years. That number is especially concerning due to the fact that 30% of all predicted species (Eukaryotes and Prokaryotes) worldwide are insects (Mora et al. 2011) and also because up to 80% of insects are still undescribed (Stork 2018). Consequently, politics became increasingly aware of the ongoing biodiversity crisis and projects such as GBOL III: Dark Taxa were funded to learn more about the hidden insect diversity (Hausmann et al. 2020). But while the extinction rate of numerous taxa is higher than ever (De Vos et al. 2015), descriptive taxonomy and morphological identification of such complex insect groups is still a rather slow process. One of the advancements in species identification and delineation, the DNA barcoding approach (Hebert et al. 2003), has helped immensely to speed up the process of species identification, the detection of new species, the evaluation of species complexes, and the interpretation of unclear systematics (Hüber et al. 2023, Goldstein and DeSalle 2011, Blagoev et al. 2009). Combining innovative methods with classic morphology is a cost and time-efficient way to tackle hidden diversity (Padial et al. 2010, Schlick-Steiner et al. 2010). Another promising new technology that keeps moving in our daily lives is advanced artificial intelligence (AI). There are many examples of how to advance biological research with those new technologies: Toscano-Miranda et al. (2022) list and compare, for instance, the applications of AI in pest control, Folliot et al. (2022) used machine learning applications in combination with acoustics to monitor pollinating insects, wood use and the ecology in a forest. Wührl et al. (2022) presented a promising state-of-the-art insect sorting device, the “DiversityScanner”, powered by a convolutional neural network (CNN). It identified specimens down to family level with a success rate of up to 100% (on average 91,4%), depending on the family they belonged to.

The better and finer scale those automated identifications become, however, the more opportunities arise for advances in insect research: one conceivable application could be only to highlight specimens that are not possible to align with a certain group the algorithm is able to recognize. Targeted evaluation without the expensive and time-consuming hand-picking would be possible (Wührl et al. 2022).

As is true for the DNA barcode system, neural networks can only be as good as the reference they are based on or trained with. As barcodes change over time (Hebert et al. 2003), depending on the data available for the clustering algorithms, neural networks can distinguish categories based on the amount and quality of the images used to train them.

Our study is based on the data of a parasitoid wasp family, the Diapriidae (Hymenoptera) obtained in the framework of the GBOL III project (Hausmann et al. 2020). Those parasitoids play important roles in the ecosystem, e.g., for pest control, and are therefore even used commercially in agriculture (e.g., *Trichopria drosophilae* to fight the invasive pest *Drosophila suzukii* (Rossi Stacconi et al. 2019)). And although those tiny (1,2–4,5mm) wasps can be found worldwide, they get little attention, and the biology of those insects is barely known (Johnson 1992). The known diversity of Diapriidae is limited to about 2000 described species, which is probably only the tip of the iceberg (pers. comm. P. Hebert). In the framework of GBOL III: Dark Taxa project, one of the two local subfamilies was further looked into as a proof of concept how to tackle highly diverse groups with disproportionately high rates of unknown diversity. The GBOL dataset is highly suitable for classification with AI because thousands of specimens were photographed, barcoded, and (therefore reliably and fine-scaled) identified, allowing a robust foundation for the network training. Our work should be interpreted as proof of concept that AI can be a valuable and fast way to evaluate extremely species-rich taxa with high levels of cryptic diversity or huge bulk samples.

## 2. Materials and Methods

### 2.1 Dataset

The dataset used for automated classification includes 11 different genera of parasitoid wasps, out of which 10 belong to the Diapriidae family and Diapriniiae subfamily. Only one taxon, the genus *Ismarus,* is of the Ismaridae family. Both the Diapriinae subfamily and the Ismaridae were picked for the proof of concept since their diversity, while still challenging, is way less incomprehensible, and the identification is less demanding as would be the case for the more diverse and abundant subfamily Belytinae. The specimens were mostly collected in southern Germany, mainly in Bavaria. Since 2011, malaise traps have been set regularly to cover various (even the most specialized) habitats, ranging from private gardens to the high alpine region. A complete list of the evaluated specimens and their location data is attached in the supplementary data. DNA barcoding was then used to identify the species.

Images of other hymenopteran species were pooled into one more class: “other Hymenoptera” comprises 121 images of other Hymenoptera like Braconidae, Ichneumonidae, Chalcidoidea, and also some Diapriidae that do not belong to the 10 previously mentioned classes since they belong to the subfamily Belytinae. The word “class” will from here on, refer to target groups that belong together and are to be sorted. It does not refer to the taxonomic, hierarchical term.

For image capture, we employed two systems: an Olympus camera E-M10 with a Novoflex Mitutoyo Plan Apo 5x microscope lens, controlled by the OM Capture software (version 3.0) was used to take deep-focused images by stacking 70-130 individual images. In addition, we took images with a prototype of the Entomoscope (Wührl et al 2023). All specimens were photographed in ethanol, mimicking the light and sample conditions used for the DiversityScanner. All images were subsequently stacked using Helicon Focus (version 8).

We used 2257 color images in our study, as summarized in Table 1. One additional test dataset has been curated to evaluate the outlier detection performance. Detailed taxonomy and the number of images in these test datasets are presented in Table 2. DNA barcoding and morphological (expert knowledge) methods were applied to identify the species. All images, including their barcoding data, are accessible on request.

**Table 1.** Taxa and the number of images used for training, validation, and testing the neural network split by sex.

| Genus | Training | Validation | Testing |
| --- | --- | --- | --- |
| *Aneurhynchus* | 104 | 11 | 20 |
| *Basalys* | 306 | 35 | 60 |
| *Coptera* | 85 | 9 | 17 |
| *Entomacis* | 71 | 8 | 14 |
| *Idiotypa* | 42 | 5 | 19 |
| *Ismarus* (Ismaridae) | 61 | 7 | 12 |
| *Monelata* | 115 | 13 | 23 |
| *Paramesius* | 110 | 13 | 22 |
| *Psilus* | 56 | 6 | 10 |
| *Spilomicrus* | 114 | 12 | 22 |
| *Trichopria* | 564 | 63 | 111 |
| Other Hymenoptera | 93 | 10 | 18 |
| female | 713 | 79 | 140 |
| male | 915 | 103 | 180 |
| unknown | 93 | 10 | 18 |
| Total | 1721 | 192 | 338 |

**Table 2.** Test dataset for outlier detection.

| Label | Descriptor | Image count |
| --- | --- | --- |
| Diapriidae, Belytinae | Parasitoid wasp | 52 |
| other insects | e.g. Aleoarthripidae: *Aleoarthrips*, Coleoptera: *Anisandrus*, Phoridae: *Megaselia*, etc. | 149 |

### 2.2 Data Preprocessing

In the computer vision field, the efficiency of model training and classification accuracy is significantly influenced by the quality and preparation of input images. This section delineates the preprocessing steps to prepare the insect image dataset for effective machine-learning model training.

#### Crop and Resize Using GroundingDINO

To enhance the model’s focus on the insect and to minimize background noise, images are first cropped to the Region of Interest (ROI) using the GroundingDINO model (Liu et al., 2023), as depicted in Figure 1. This model employs a zero-shot object detection approach, leveraging image and text features to predict bounding boxes around the insect based on the provided text prompt ’Insect. Wasp. Wings.’ with a box threshold of 0.29 and a text threshold of 0.25. These cropped images are then resized to a uniform size of 224x224 pixels. This standardization step preserves critical insect features for further processing.

**Figure 1.**
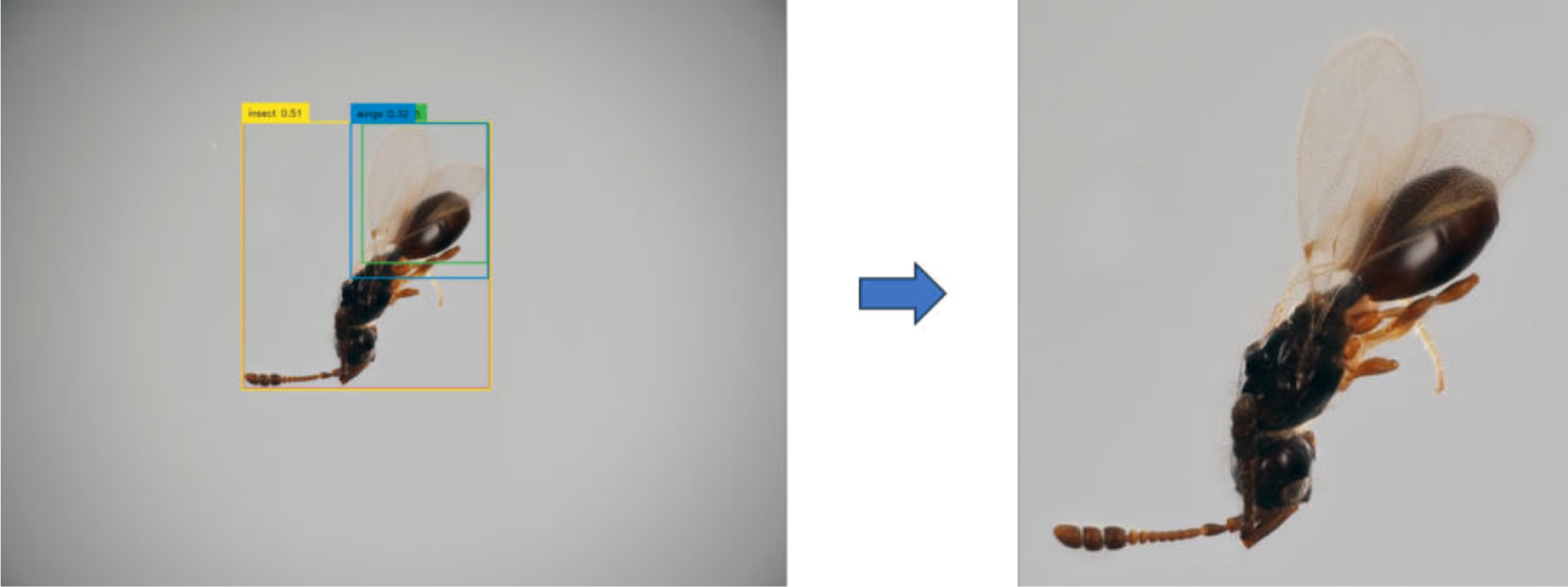
Object Detection using GroundingDINO with subsequent Cropping visualized

#### Data Augmentation

To enrich the dataset and prevent overfitting, data augmentation techniques such as horizontal and vertical flip, rotation (-30° - +30°), horizontal shift (1-8% of the image width), vertical shift (1-8% of the image height) and zooming in or out (up to 8%) are applied. These techniques help the model learn from a more diverse representation of insect features.

#### Final Dataset Compilation

The preprocessed images are compiled into the final dataset and randomly split into a training dataset (∼69%), a validation dataset (∼11%), and a testing dataset (∼20%), considering class imbalance to effectively assess the model’s performance and generalizability. These steps ensure the dataset is thoroughly prepared for the subsequent model training and evaluation phases, establishing a solid foundation for precise and robust insect classification.

### 2.3 Deep Learning Model Architectures

Three different deep learning models were selected and evaluated in this study: ConvNeXt, BEiTv2, and YOLOv8. These models were selected for their proficiency in handling complex computer vision tasks, particularly in identification and classification. Our approach is grounded in transfer learning and fine-tuning methodologies, ensuring the models are adapted to our specific requirements.

#### ConvNeXt XLarge

(Li et al. 2022) is an advanced Convolutional Neural Networks (CNNs) variant known for its exceptional feature extraction capabilities. It incorporates multiple layers designed to process and interpret intricate image details, leveraging advanced activation functions and optimizers to ensure efficient learning and high classification accuracy. The model supports multi-label classification with a sigmoid activation function and handles an image size of 224 x 224 pixels with a batch size of 32 and stochastic depth regularization with a rate of 0.3. The class weights are similarly adjusted for genera classes.

The second model is **BEiTv2** (Peng et al. 2022), a Transformer-based model adapted to understanding and interpreting complex image patterns. Its unique attention mechanism is instrumental in identifying subtle variations within images, making it a crucial tool for ensuring model stability and robustness under diverse imaging conditions. The model processes images of 224 x 224 pixels with a batch size of 32 and employs a dropout regularization of 0.3 applied to Attention-MLP (multilayer perceptron) blocks. Class weights for genera classes are weighted by a factor of 3.

The third model is **YOLOv8** (Jocher et al., 2023), the latest iteration in the YOLO (You Only Look Once) series, chosen for its rapid object detection capabilities, which are also well-suited for classification tasks. Its architecture, balanced for speed and accuracy, makes it ideal for real-time applications where immediate and precise classification is essential. Due to the framework’s limitation in not supporting multi-label classification, we train two separate models – one for genus classification and one for sex determination. Both models leverage ImageNet pre-training weights (Russakovsky et al. 2015), and all layers are made trainable, an approach that maximizes learning from our dataset. These models are designed for multi-output classification, utilizing a softmax activation function. They are capable of processing larger images of 640 x 640 pixels, operating with a batch size of 64 and incorporating a dropout regularization of 0.3. The class weights for both models are set to default. This configuration ensures optimal performance and accuracy in our classification tasks. In conclusion, the architecture of each model has been tailored to meet the specific requirements of this project. ConvNeXt’s advanced convolutional approach, BEiTv2’s attention-based mechanism, and YOLOv8’s speed and precision collectively contribute to the successful implementation of the classification tasks in this study.

### 2.4 Training Setup and Process

For classification, a standard personal computer with a powerful NVIDIA RTX 4080 GPU was used with Python (3.10), TensorFlow (2.10.1), PyTorch (2.0.1), Keras, CUDA (11.7), and Anaconda software. This integrated environment provides the efficiency and flexibility to train deep learning models. In the training process, all three models were trained for a maximum of 150 epochs using the AdamW optimizer with a consistent learning rate of 0.001. We employed a 4-fold cross-validation approach to optimize model performance, This allowed us to assess the models’ performance on different subsets of the data, mitigating the risk of overfitting and providing a more robust evaluation of their generalization capabilities. In addition to cross-validation, we also applied early stopping, model checkpointing, and learning rate reduction techniques, with training progress monitored. Notably, model weights were saved whenever improvements were observed during validation. BEiTv2 and YOLOv8 utilized categorical cross entropy for loss functions, while ConvNeXt employed binary cross entropy.

### 2.5 Outlier Detection

An algorithm for automatic classification is expected to reliably differentiate between insects that belong to the predefined classes for classification and between specimens that do not belong to these classes. This allows the automatic filtering of collections that have not previously been pre-sorted for the predefined classes. For this reason, the algorithm first classifies a specimen into one of the two groups, “Hymenoptera for classification” and “Non-Hymenoptera”. This pre-filtering is carried out by a one-class Support Vector Machine (OCSVM) based on the BEiTV2 - a pre-trained deep learning model with ImageNet weights. Puls et al. (2023) have demonstrated that ViTs perform best for this task. During this process, the classification layer is removed, leaving the model to serve as an effective feature extractor. This model transforms the input images into a lower-dimensional feature space, capturing low-level and high-level image features. Subsequently, a one-class Support Vector Machine (SVM) on these feature representations extracted from the training dataset is trained. Any new testing data that falls within the boundary of the one-class SVM is assigned to the trained class, and data points outside the boundary are declared as outliers or Non_Hymenoptera.

Principal Component Analysis (PCA) is then employed to reduce the dimensionality of the data from 1024 to 128 features per image to maintain data quality while reducing computational complexity. In this next step, the data is normalized using the mean and variance of the training data set. Subsequently, the one-class SVM is trained on the reduced and normalized feature representations. Since this approach doesn’t involve training a neural network, there’s no need for a separate validation dataset. Instead, the validation dataset is combined with the training dataset for training the one-class SVM on the positive class, making it suitable for detecting outliers, which, in this context, are the other insects.

The entire approach is implemented using the open-source machine learning library Scikit-learn (Pedregosa et al. 2011). Parameter tuning is performed through a grid search to optimize the OCSVM’s performance.

## 3. Results

### 3.1 Classification performance metrics

The performance metrics for genus and sex classification of the three different deep learning (DL) models are given in Table 3. The performance metrics include the classification accuracy and the F1-score (a metric that combines precision and recall) across four training runs using 4-fold cross-validation.

**Table 3.** Performance metrics of three different deep learning architectures for genus and sex classification. A value of 1 corresponds to 100%.

| Architectures | Genus Accuracy | Genus F1-score | Sex Accuracy | Sex F1-score |
| --- | --- | --- | --- | --- |
| **BEiTv2** | 0.96 | 0.95 | 0.97 | 0.98 |
| **ConvNeXt XLarge** | 0.94 | 0.94 | 0.95 | 0.96 |
| **YOLOv8** | 0.89 | 0.90 | 0.94 | 0.94 |

The performance metrics show that BEiTv2 consistently outperforms the other models in genus and sex classification tasks. ConvNeXt XLarge also exhibits strong performance, while YOLOv8 performs competitively, albeit with a reduction in accuracy and F1-score compared to the two other models. For this reason, only the classification results of the best-performing model, BEiTv2, are presented below.

The classification results for the 11 predefined classes of Hymenoptera and one “other-Hymenoptera” class are depicted in a confusion matrix in Figure 2 and Figure 3 for the tasks of genus and sex classification.

**Figure 2.**
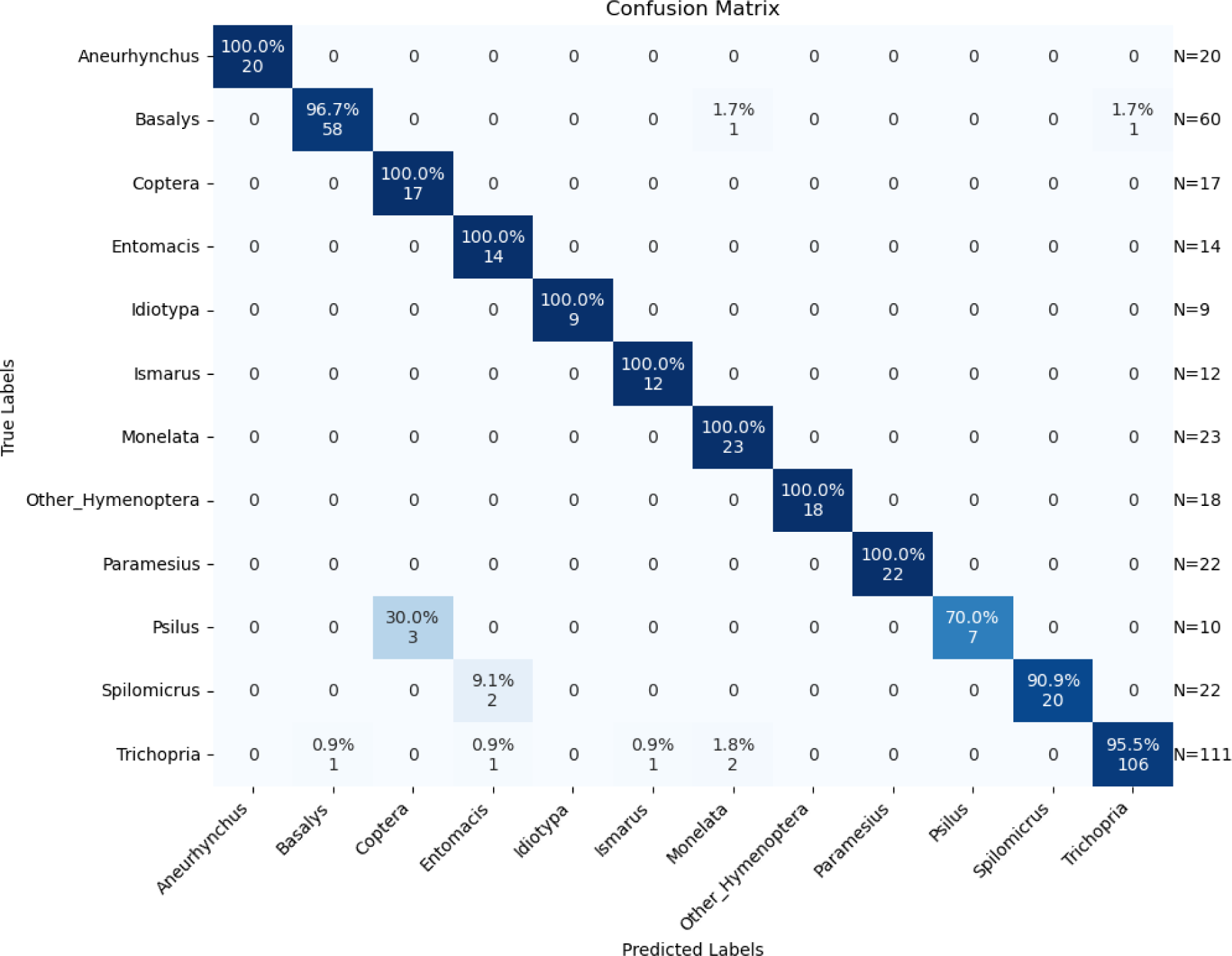
Confusion matrix with genus classification results of the BeitV2 model.

**Figure 3.**
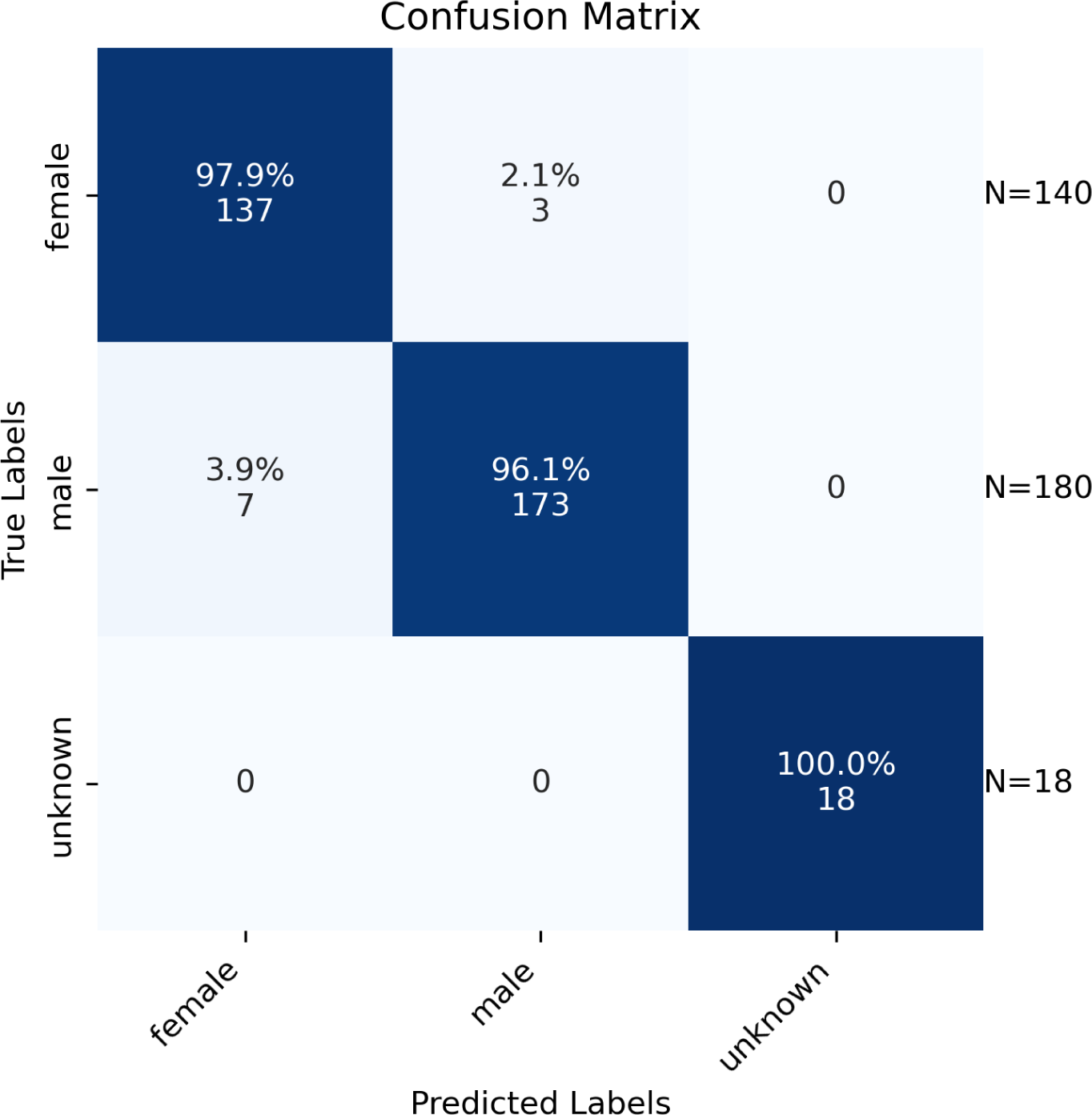
Confusion matrix with sex classification results of the BeitV2 model.

In addition, the graphs of the classification results for training and validation accuracy and loss for the BEiTv2 model are given in Figures 4, 5, and 6. These figures are for the first fold of the cross-validation training process. They provide a comprehensive view of the model’s learning progress throughout training, shedding light on its overall performance and convergence behavior.

**Figure 4.**
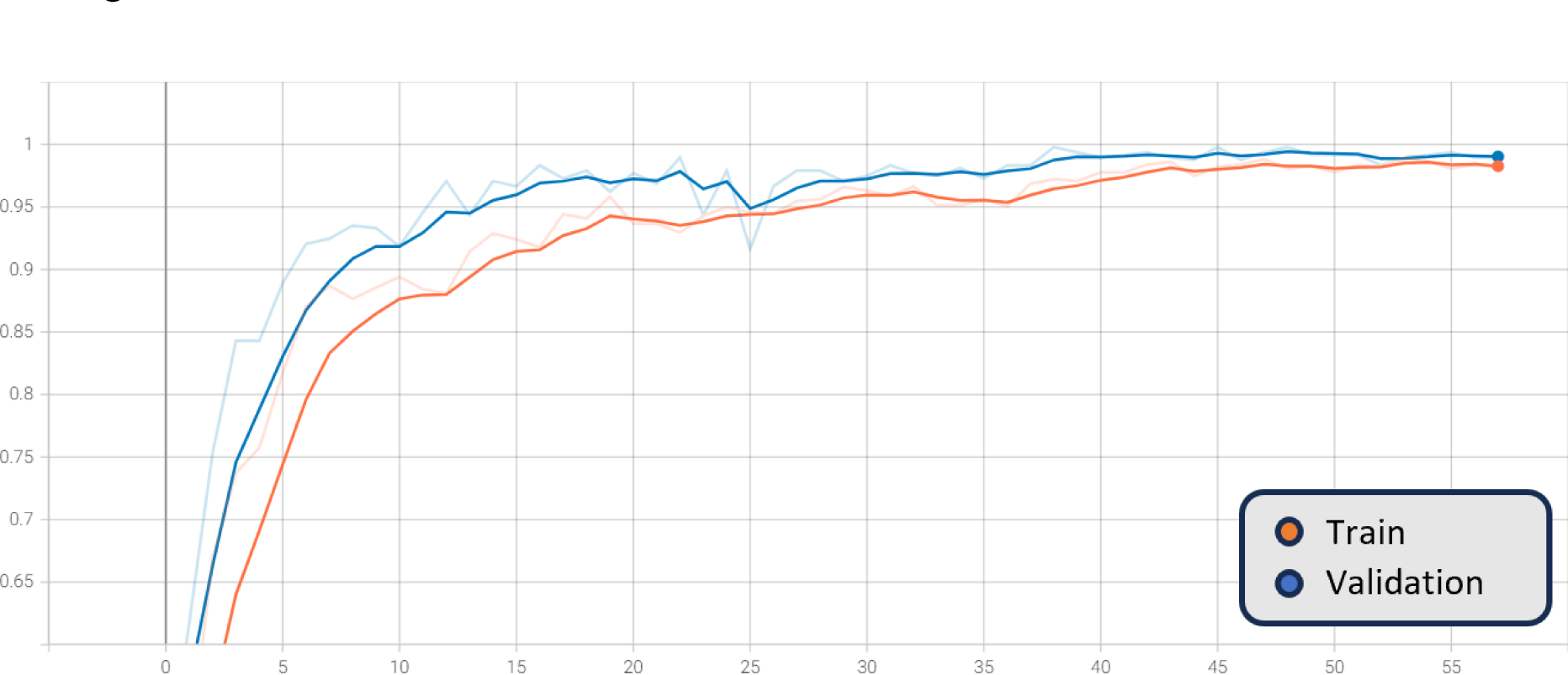
Smoothed training (orange) and validation (blue) genus accuracy, BEiTV2, with the original graph in transparent, X-axis: epoch number, Y-axis: genus accuracy

**Figure 5.**
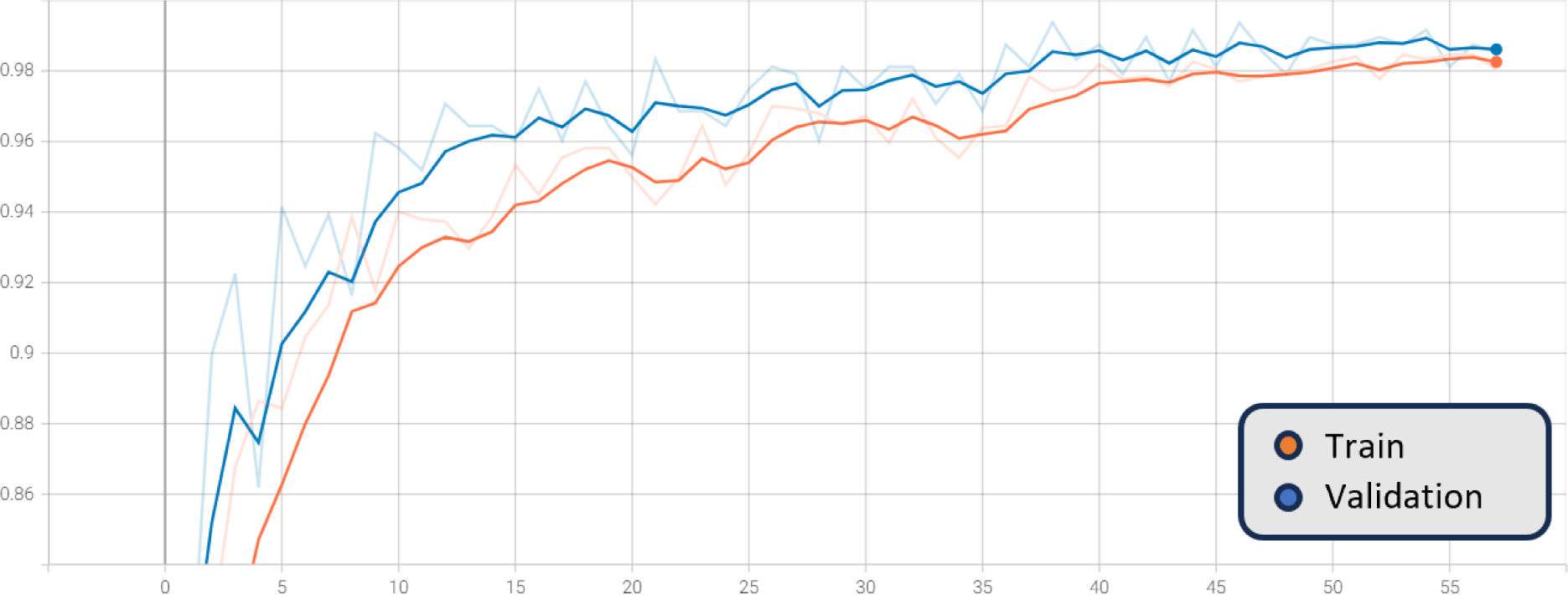
Smoothed training (orange) and validation (blue) sex accuracy, BEiTV2, with the original graph in transparent, X-axis: epoch number, Y-axis: sex accuracy

**Figure 6.**
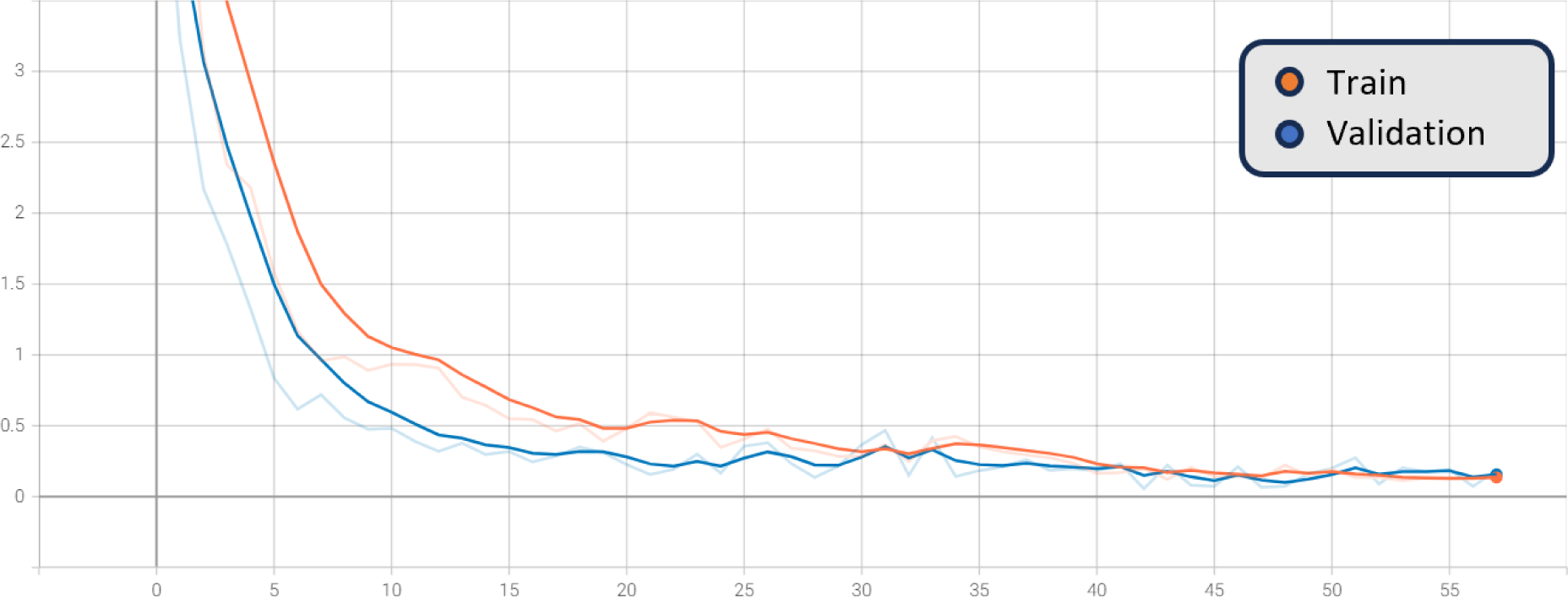
Smoothed training (orange) and validation (blue) combined genus and sex loss, BEiTV2, with the original graph in transparent, X-axis: epoch number, Y-axis: loss

The figures show a steady increase in accuracy and a corresponding decrease in loss, suggesting the model is learning effectively. Notably, the close alignment of the training and validation curves indicates that the model is not overfitting, as it performs similarly on both seen and unseen data. Moreover, the absence of a plateau in improvement or a significant gap between training and validation performance suggests that underfitting is not occurring. Hence, the model exhibits a balanced learning trajectory, which suggests robustness and reliability when applied to similar unseen data.

### 3.2 Class Activation Maps

Class Activation Mapping (CAM) (Zhou et al. 2016) is a technique for generating heat maps to highlight class-specific regions of images that impact the classification result. In Figure 7, heat maps for two different insect specimens are given as examples: the genus *Paramesius* (top) and *Spilomicrus* (bottom). The left side represents heat maps associated with the predicted genus. The antennae, the head, and the thorax are consistently significant in predicting the genus. On the right side, the heat maps related to sex prediction are displayed, where the antennae are crucial for sex prediction. These results show that the classification algorithm considers features similar to those of a taxonomic expert.

**Figure 7.**
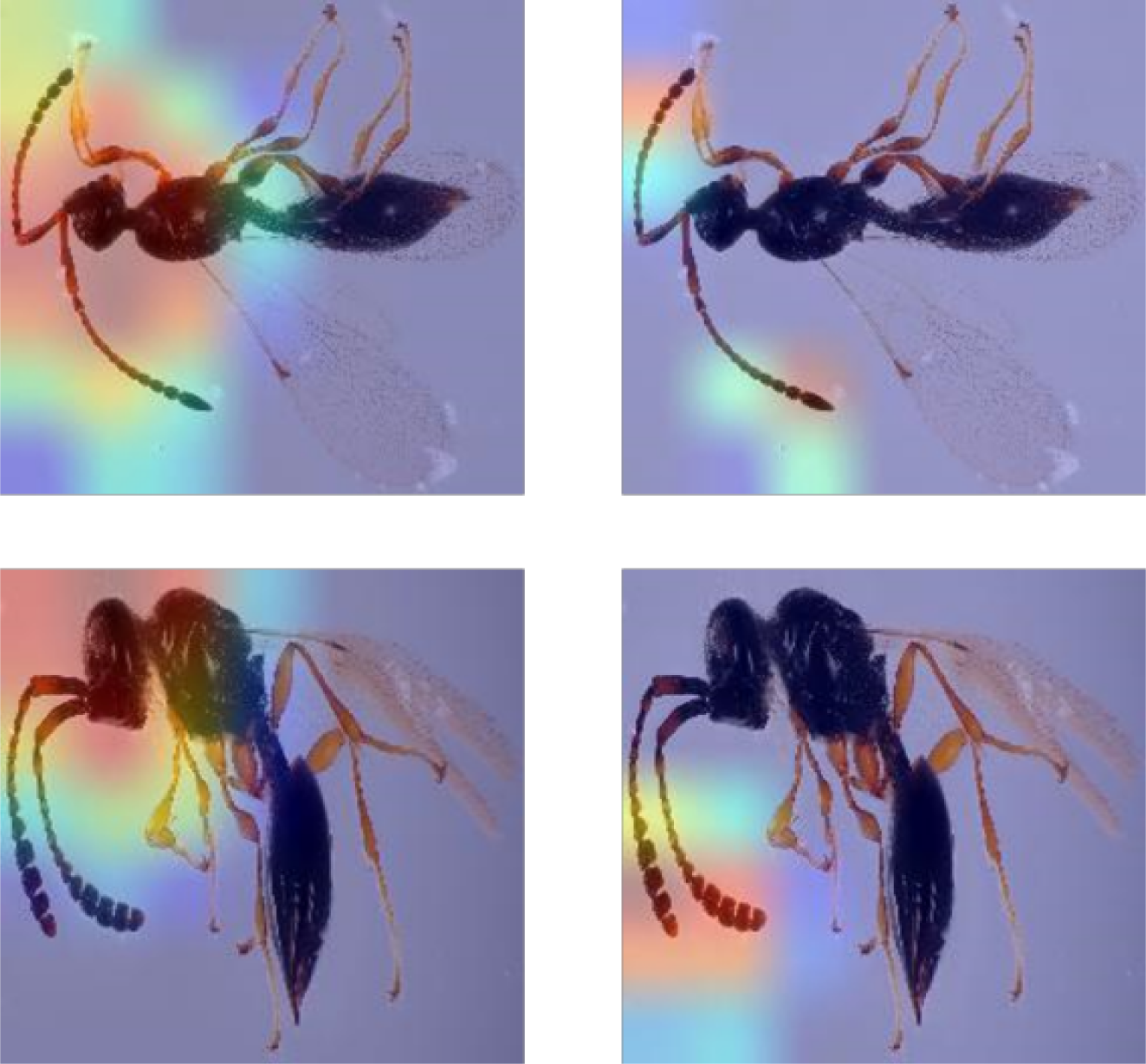
Class activation heatmaps for genus classification (left side) and sex classification (right side). Red areas indicate regions with higher weighting in the classification.

### 3.3 Outlier Detection

The outlier detection method was assessed using two different test datasets: one is described in Table 2, and the other is a split of our main dataset. The results are visualized in Figure 7. Out of 652 images in total, the method misclassified 23 of them.

Notably, this approach achieved 100% accuracy on the test split of our dataset, which was expected since our outlier model was trained specifically on this dataset. Regarding the “Other Insects” category, this model demonstrated its prowess by identifying 90.6% of the images as outliers. This indicates the model’s ability to distinguish these insects from Hymenoptera effectively. The Diapriidae Belytinae dataset, as part of “Other Hymenoptera,” presented a unique challenge. It included variations in image quality, background, and differences in camera sources. Despite these challenges, our model detected 82.7% of the images as inliers, underscoring its potential for accurate classification even under adverse conditions.

**Figure 7.**
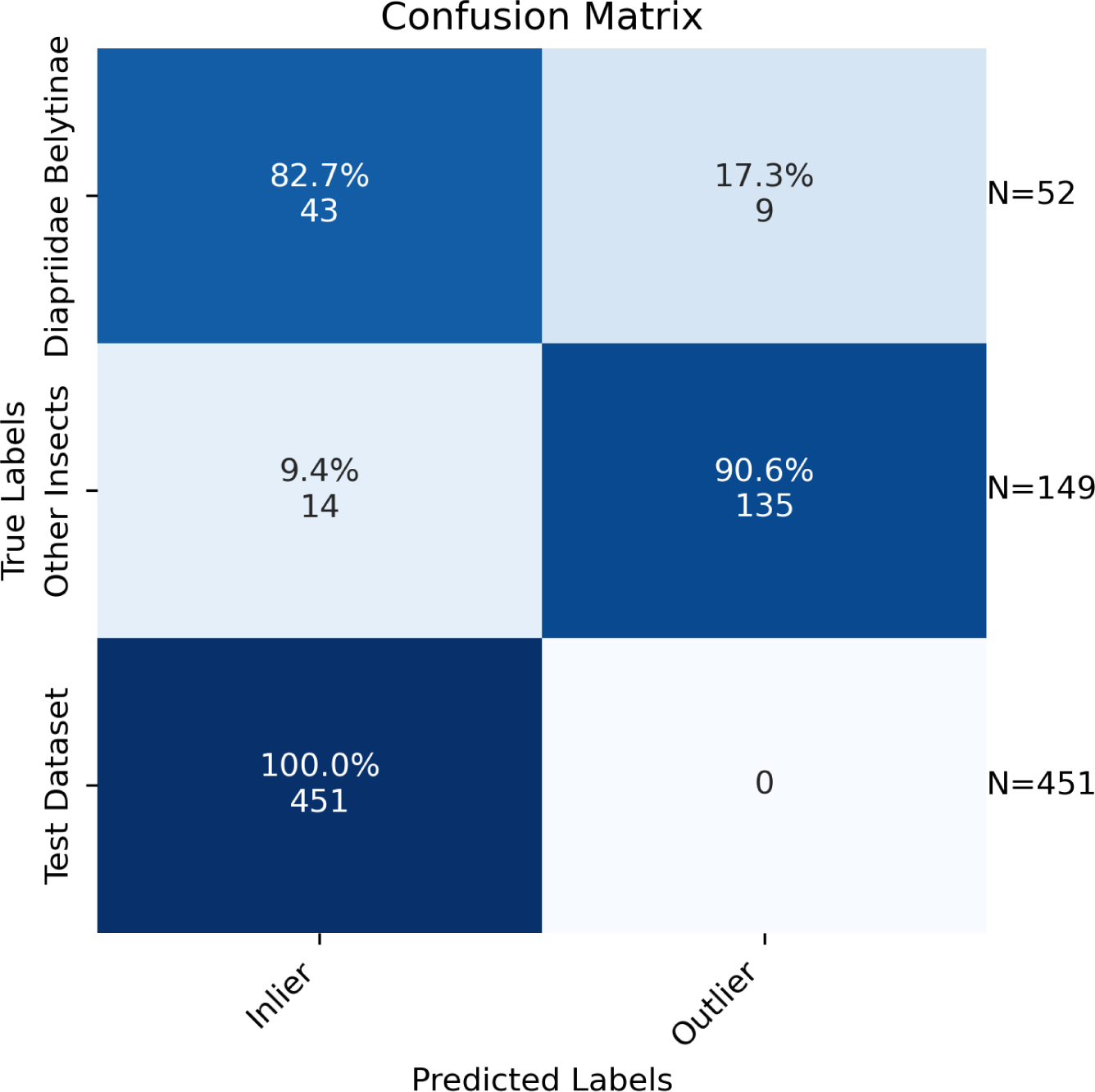
Confusion Matrix for the Outlier Detection

## 4. Discussion

The shown network approach is restricted to the European Diapriidae fauna, particularly to the subfamily Diapriinae. The reason for that is that within the framework of the GBOL III project, specimens/species of this specific subfamily were investigated and barcoded as proof of concept. The Diapriidae (even if the dataset is limited to German material only) are just too diverse and complex to be investigated in such a short period of time. Nevertheless, most of the genera that were subject to our approach are worldwide distributed. Also, there are many species, e.g., *Spilomicrus formosus*, that even inhabit several continents (in this case, Europe, Asia, and North America), which makes our DL model a powerful tool regarding the fact that over 90% of the sampling area of the barcoded material was limited to Bavaria (Germany).

The success rate at which the DL model was able to distinguish between different genera was really high (up to 100%). Exceptions could be detected distinguishing between the genera *Psilus* and *Coptera.* Having a closer look into those was not surprising. Both genera are closely related and look very much alike. Although the genus *Psilus* was described by Panzer in 1801 and *Coptera* by Say (1836) 40 years ago, there was still confusion on how to tell them apart (Nixon 1980). The most reliable morphological feature is the wing. *Coptera* has it folded lengthwise, while *Psilus* does not have that fold. But since both genera usually lay on their side with applied wings, it is merely impossible to distinguish between them without changing their position to a dorsal view. Another obstacle we faced was that there was not enough material to train the models on rare taxa. *Idiotypa, Diapria,* or *Tetramopria* are genera with low species and individual counts.

The class activation heatmaps highlight, as expected, the antennae of the insects, which taxonomists also use to distinguish between sexes. What was less expected was that the CAMs highlighted the head region. Although the head shape could be used to identify genera, other body features would be used by a specialist: the wing venation (which is often not visible in the images) and the shape of the abdomen (which is not always helpful and dependent on its orientation) would be in our example in fig. 7 much more intuitive to distinguish *Paramesius* and *Spilomicrus.* Therefore, CAMs might have the potential to find descriptive characters for species descriptions in the future.

But while the algorithm can not identify them down to genus level, it can determine their family and therefore be used to sort specifically for rare, unidentifiable specimens, which would save even a specialist vast amounts of time due to the generally high specimen numbers of most diaprids.

In furthering this research, we developed a web application to make the identification process more accessible and user-friendly (Both et al., 2023). However, it is crucial to note that the application’s accuracy is highly dependent on the quality of the images used. Only high-quality lab images with consistent and comparable illumination are suitable for the app’s analysis. Images taken with a smartphone, which often vary in quality and lighting conditions, are unlikely to yield reliable results. This limitation emphasizes the need for standardized image-capturing methods to ensure the app’s effectiveness in species identification.

## Conclusion

AI has been proven to be a reliable and efficient tool to identify the highly diverse taxon Diapriinae down to the genus level in Europe. One of the greatest advantages lies in the fact that a user does not have to have a profound knowledge of morphology or other taxonomic experience that took specialists great effort and time to acquire. Making those groups available for completely different research fields, such as ecology or pest control, is a great advancement and a cheap and non-invasive alternative to (meta-) barcoding-based species identification. This technology has and should be further developed and can be applied to all sorts of species groups, e.g., other parasitoid wasps. Another potential application could be to power the DiversityScanner with those new DL models to allow more accurate delimitations and targeted specimen picking.

## Acknowledgments

We also want to thank the students Viktor Deines and Jerome Anton for putting in countless hours to image all the investigated specimens.

Our work is part of the German Barcode of Life III: Dark Taxa project and was funded partially by the German Federal Ministry of Education and Research (FKZ 16LI1901B). It was also supported by funding from the Museum für Naturkunde Berlin and the program Natural, Artificial and Cognitive Information Processing (NACIP) of the Helmholtz Association.

